# Powassan virus infects mouse testes and reveals a protective role for phagocytic cells

**DOI:** 10.64898/2026.07.17.739246

**Authors:** E. Eldridge Hager-Soto, Esteban Arroyave, Eduardo Eyzaguirre, Omar A. Saldarriaga, Alexander N. Freiberg, Shannan L. Rossi

## Abstract

Powassan virus (POWV) is an emerging tick-borne *orthoflavivirus* with consistently increasing annual prevalence. Though understood as a vector-transmitted virus associated with neurological disease, a 2016 fatal case of an immune-compromised 63-year-old man reported testicular pain with postmortem detection of viral antigen in the testes suggested an altered tropism. The extent to which POWV can infect testes remains unclear. We used a BALB/c mouse model of POWV to elucidate the potential of POWV to establish infection in testes and model a potential mechanism of testicular tropism. Here, we report that POWV productively infects the male reproductive tract of mice. We find that POWV consistently infects mouse testes and epididymides at 3-and 5-days post-infection (dpi) with peak titers measuring 6.1 log_10_ PFU/gram tissue in testes. Furthermore, we find that at 5dpi POWV-infected mice testes are significantly higher in mass than uninfected controls. Due to macrophages being implicated in driving testicular pathogenesis in Zika virus, a related *orthoflavivirus*, we depleted phagocytic cells using clodronate liposomes prior to POWV infection and compared to control liposome treated mice. Depletion of phagocytic cells was associated with higher and sustained viremia, and significantly higher viral organ burden in the male reproductive tract of mice. Gene expression analysis revealed a robust antiviral, inflammatory, and chemokine response in the testes that was further amplified following phagocytic-cell depletion. These findings implicate the male reproductive tract as a previously underrecognized target of POWV disease and identify phagocytic cells as key in limiting viral infection of testes in mice.

## 1. Introduction

Powassan virus (POWV) is an emerging tick-borne orthoflavivirus endemic to the northeastern United States and far-East Russia^1,2^. Disease associated with POWV infection in people initially presents as fever, headache, and vomiting, and severe disease can result from neuroinvasive infection leading to encephalitis and/or meningitis^2^. Neuroinvasive POWV infection has a mortality rate of approximately 10%, and patients who recover from acute disease struggle with neurological sequelae in around 50% of cases^3,4^. POWV disease currently has no cure or treatment, and no approved vaccine exists for prevention. POWV remains a rarely diagnosed disease; however, 2025 marked the fourth consecutive year with the highest reported POWV cases at 63 reported infections^5^.

Although POWV typically presents with non-specific or neurological symptoms, a lethal case described in 2016 initially reported testicular pain^6^. Testicular ultrasonography revealed orchiepididymitis, and post-mortem analysis detected POWV antigen in the patient’s testis. The patient had been previously diagnosed with follicular lymphoma and was receiving rituximab. Due to the paucity of POWV infections and autopsy reports following death, the incidence of even one confirmed case of testicular infection is noteworthy and merits further investigation.

The potential for POWV to establish testicular infection is not without precedence; viral infection of the testes has been well documented for a variety of pathogens across different viral families^7^. Mumps and human immunodeficiency viruses are classical examples, and filoviruses like Ebola and Marburg viruses are now recognized as testicular tropic, especially after mounting epidemiological evidence emerged from the 2014-2016 West African epidemic^8–12^. Zika virus (ZIKV), another arthropod-borne orthoflavivirus, has been shown to infect the male genital tract, including testes^13^. Epidemiological evidence as early as 2008 suggested the possibility of sexual transmission, and therefore testicular infection; however, these observations remained largely unexplored until the American ZIKV epidemic almost a decade later^14,15^. Recently, Oropouche virus was detected in a traveler’s semen in 2024, and severe fever with thrombocytopenia syndrome virus has been found to infect the male reproductive tract, causing orchitis, impaired spermatogenesis, and seminal viral shedding^16,17^. These cases of testicular tropic viruses underscore the need for more dedicated research into viral testicular infection, particularly when incidental evidence emerges.

Based upon evidence from arboviral infections and the aforementioned clinical POWV case, we sought to investigate the testicular tropism of POWV infection. Previous literature documenting testicular infection of POWV in mice has been incidental and inconsistent. One 2020 study found in 65% of tested male mice between 11-55 focus forming units (FFU) equivalents/μg RNA in animals that had succumbed to infection, and a report on POWV tick transmission in mice in 2022 reported one male mouse with between 10^5^-10^6^ FFU equivalents/ug RNA at 28 days post-infection^18,19^. Furthermore, testicular tropism was supported only by detection of viral RNA and limited to study endpoints. Our study seeks to fill these gaps by systematically quantifying the dynamics and potential mechanisms underlying POWV testicular tropism.

Using the previously established BALB/c murine model for POWV, we found that mice consistently harbor infectious virus during acute infection in testes and epididymides. We report that infected mice testes significantly increase in mass day 3-5 post-infection but show no signs of histopathology, and viral antigen localizes within the seminiferous tubules. Furthermore, we aimed to evaluate the role macrophages play in controlling POWV at early stages of infection as a cell type susceptible to POWV infection^20^. As macrophages have been implicated in delivering ZIKV to the testes following a “Trojan horse”-like mechanism, we were interested in determining if POWV may follow a similar mechanism should it display a tropism for the testes^21^. We found that following phagocytic cell depletion, POWV viremia significantly increases and is prolonged, and testes and epididymides had significantly higher viral loads. Gene expression analysis showed induction of an interferon driven pro-inflammatory response in the testes of POWV-infected mice, with phagocytic-cell depleted mice showing heightened levels of pro-inflammatory transcriptional activation. These data collectively inform us that POWV productively infects the mouse reproductive tract and that phagocytic cells serve as important regulators of early viral replication and inflammatory response.

## 2. Materials and Methods

### Cells and viruses

Vero CCL-81 cells (American Type Culture Collection) were grown in Dulbecco’s minimal essential media (DMEM, Gibco, Thermo Fisher Scientific, Waltham, MA) with 10% fetal bovine serum (FBS), 200 U/mL penicillin and 200 mg/mL streptomycin (Gibco, Thermo Fisher Scientific, Waltham, MA). Human primary Sertoli cells (HSerCs) (IX Cells Biotechnologies) were maintained in Sertoli cell basal media (IX Cells Biotechnologies) supplemented with 10% FBS and supplemented with Sertoli cell antibiotics and growth supplement according to the manufacturer’s recommendations (IX Cells Biotechnologies). BHKsa cells were grown in Minimal Essential Medium Alpha (1X) + GlutaMAX (Gibco, Thermo Fisher Scientific, Waltham, MA) with 10% FBS, 200 U/mL penicillin and 200 mg/mL streptomycin (Gibco, Thermo Fisher Scientific, Waltham, MA). HSerCs were used within 3-5 passages after acquisition. BHKsa cells and Vero CCL-81 cells were used between 5-15 passages after acquisition.

POWV Lineage I (LB) and POWV Lineage II (Deer Tick Virus, DTV) were received from the World Reference Center for Emerging Viruses and Arboviruses (WRCEVA), University of Texas Medical Branch (UTMB, Galveston, TX). POWV LB was passaged twice in LLC-MK-2 cells and once in Vero CCL-81 cells (American Type Culture Collection, Manassas, VA) and DTV was passaged thrice in suckling mouse brain. All infectious work was conducted at BSL-3 at UTMB.

### Ethics for studies with infectious agents

Animal experiments involving infectious POWV and DTV were conducted at animal biosafety level 3 (ABSL-3) at UTMB under IACUC protocol 2507034 and performed in accredited facilities, ensuring compliance with the guidelines of the Association for Assessment and Accreditation of Laboratory Animal Care International (AAALAC).

### Virus titration by plaque assays

Plaque forming units (PFU) were determined by plaque assays on monolayers of BHKsa cells for POWV and Vero CCL-81 cells for DTV in 12-well plates^22^. Briefly, samples were diluted serially and used to infect Vero CCL-81 or BHKsa monolayers. After 1 h of adsorption with gentle rocking at 37 °C, cells were overlayed with 1XMEM (Gibco, Thermo Fisher Scientific, Waltham, MA) containing 0.8% tragacanth (Sigma Aldrich). Plates were incubated at 37 °C with 5% CO_2_ for approximately 3 days for POWV or 6 days for DTV. Monolayers were fixed using a 10% formaldehyde solution. The cells were treated with a solution of 0.2% crystal violet in 30% methanol to visualize the plaques. Data are shown as log_10_ PFU/ml, with the limit of detection (LOD) for each assay indicated on each graph by a dotted line.

Organs were collected in pre-weighed tubes to determine organ mass. Tubes were homogenized at a frequency of 30 Hz for 2 minutes prior to centrifugation at 6,000 x g for 5 minutes. Supernatants were collected and titers were determined as described above. Data are shown as log_10_ PFU/gram of tissue.

### Viral growth curve

Cells were seeded in 12-well plates and allowed to reach 80-90% confluency prior to infecting. Cells were infected at various multiplicities of infection (MOI) in a total infection volume of 100μL and incubated at 37 °C with 5% CO_2_ for 60 minutes. Cells were rocked every 15 minutes to prevent cells from drying and ensuring the entire monolayer is exposed to the inoculum. After infection, each well was washed with 1 mL of 1X PBS three times, before 1 mL cell growth medium was added to each well. At a consistent time each day post infection (dpi), 100μL of supernatant from each well was removed for freezing at −80 °C for later titration and replaced with 100μL of cell growth media. Cell monolayers were visually assessed for signs of cytopathic effect such as cell rounding, cell detachment, formation of syncytia, and cell lysis using a phase contrast light microscope.

### Animal manipulations

Male 9-10-week-old BALB/c mice were obtained from Charles River (Charles River Laboratories, Bar Harbor, ME). Groups of 5 mice were infected by intraperitoneal (IP) or intradermal (ID) route with a target dose of 100 log_10_ PFU of POWV or DTV. Uninfected control mice (n=3 per group per study) were injected with sterile 0.9% sodium chloride (Hospira, Lake Forest, IL) via the IP or ID route. After infection, mice were monitored and weighed at least once daily. Blood was collected at 1-, 2-, and 3-days post-infection by the retroorbital route. Animals were anesthetized with isoflurane for each manipulation.

For macrophage depletion studies, mice received IP injections containing clodronate-containing or control liposomes (Liposoma, Amsterdam, Netherlands) 100μL reagent/10g mouse weight 48 and 24 hours prior to infection, according to manufacturer’s recommendations. When required, perfusion was performed with PBS following euthanasia. Perfusions were performed for mice infected with DTV and in clodronate-treated and control liposome-treated mice. These mice were perfused since we anticipated viremia before 5dpi as well as during phagocytic-cell depletion. POWV-infected mice at 3dpi were also perfused. POWV-infected mice at 5dpi were non perfused.

### Flow cytometry

Macrophage populations in mouse spleen and testes were identified and quantified by flow cytometry. Individual organs were macerated through 70-µm cell strainers and digested with 0.05% collagenase type IV (Thermo Fisher Scientific, Waltham, MA) in Roswell Park Memorial Institute medium 1640 (RPMI 1640) medium for 30 mins at 37 °C and again passed through 70-µm cell strainers to prepare single-cell suspension. Red blood cells were lysed through a 3 second wash in pure molecular grade followed by addition of 10X PBS to restore isotonic conditions. Live/Dead antibody (Zombie NIR Dye, Biolegend, San Diego, CA) was incubated at 1:5000 at room temperature for 15 minutes. Cells were then washed in buffer (containing 2% FBS in sterile 1X PBS). The fluorochrome-labeled antibodies used were purchased from Thermo Fisher Scientific, Tonbo Biosciences (San Diego, CA), or Biolegend as below: violetFluor 450 anti-CD8 (Clone 53-6.7), PerCP-Cy5.5 anti-CD4 (clone RM4-5), FITC anti-CD4 (Clone GK1.5), PerCP-Cy5.5 anti-CD11b (Clone M1/70), BV421 anti-F4/80 (Clone T45-2342), and APC anti-Ly6G (Clone 1A8). Cells were fixed in paraformaldehyde 2% for 16 h at 4 °C. Cell samples were processed on a BD LSR Fortessa and data were analyzed using FlowJo 10 (BD).

### Histology

Tissue samples were fixed in a solution of 10% neutral buffered formalin (Thermo Fisher Scientific, Waltham, MA) for 24 hours, then exchanged with new formalin and fixed for 31 days. Formalin solution was replaced with 95% ethanol and sent to UTMB’s Anatomic Pathology Laboratory core facility, where organ samples were embedded in paraffin and cut into 5µm sections. Representative slides from each animal were stained with hematoxylin-eosin (H&E) or left unstained for use in immunofluorescence analysis. A blinded board-certified pathologist interpreted the H&E slides. Images were captured using brightfield channel of inverted fluorescence microscope (Olympus-IX73) and Olympus cellSense Standard 2.3 software.

### Immunofluorescence assay

Slides were prepared for immunofluorescence by heating 5µm sections of paraffin embedded mouse testes at 65 °C overnight for dewaxing, followed by incubations in multiple xylene baths and a series of graded ethanol (Millipore, Sigma, Burmington, MA). Antigen retrieval was performed through a 30-minute incubation in 90 °C citrate buffer pH 6 (Abcam, Cambridge, UK) diluted in PBS (Gibco, ThermoFisher Scientific, Waltham, MA). Sections were washed in deionized water and incubated in a blocking buffer (5% normal goat serum in PBS) for 15 min at 37 °C in a humidified chamber. Primary antibody was incubated for 16 hours in a humidified chamber at 4 °C with 4G2 antibody targeting flavivirus envelope protein (Abcam, Cambridge, UK) at 1:200 dilution and DDX4 antibody targeting migratory germ cells (Abcam, Cambridge, UK) at 1:500 in blocking buffer. Samples were washed in PBS and incubated with 0.2% Sudan black B (SigmaAldrich, Burlington, MA) diluted in 70% ethanol for 12 min at 37 °C in a humidified chamber. Samples were then stained for 45 min at 37 °C with a secondary antibody in a humidified chamber at a dilution of 1:1000, AlexaFluor 594 anti-mouse IgG (Thermo Fisher Scientific, Waltham, MA) or AlexaFluor 488 anti-rabbit IgG (ThermoFisher Scientific, Waltham, MA). The slides were incubated with DAPI for 5 minutes at room temperature followed by five PBS washes prior to mouting with Vectashield (Vector Labs, Newark CA). Images were captured using an inverted fluorescence microscope (Olympus-IX73) and Olympus cellSense Standard 2.3 software.

### Gene expression analysis

RNA from mouse testes was extracted using phenol-chloroform extraction. Clarified supernatant from organ homogenate inactivated in TRIzol (Thermo Fisher Scientific, Waltham, MA) was incubated with 1 part chloroform (SigmaAldrich, Burlington, MA) to 4 parts sample and incubated for 3 minutes. Tubes were centrifuged at 12,000 x g for 15 minutes, and upper aqueous phase was removed to 0.5 mL isopropanol (SigmaAldrich, Burlington, MA) for 10-minute incubation. Samples were centrifuged for 20 minutes at 20,000 x g and supernatant was replaced with 75% ethanol (SigmaAldrich, Burlington, MA). Samples were shaken and re-centrifuged. Supernatant was discarded, and RNA was eluted in RNase-free molecular grade water (Invitrogen, Waltham, MA). All samples were diluted to 30 ng/µL prior to hybridization. RNA quality and concentration were confirmed using Nanodrop spectrophotometer (DeNovis DS-11+ Spectrophotometer). Gene expression analysis was performed using a NanoString nCounter SPRINT profiler using NCounter Mouse Myeloid V2 and NCounter Mouse Immunology V2 panels (Bruker Spatial Biology). Analysis was performed using Mouse Myeloid panel supplemented with non-overlapping genes from Mouse Immunology panel. Data analysis was performed using nSolver (nSolver Analysis Software version 4.0) and Prism 11 (GraphPad Prism version 11.0.0) software. Data normalization was performed on raw gene counts by the nSolver software using housekeeping genes and positive controls.

### Quantification and statistical analysis

For viral organ load and viremia, data were compared using a one-way ANOVA when comparing between multiple groups or an unpaired student’s t-test when only comparing two groups. Viral titers were log_10_ transformed before statistical tests were applied. Volcano plots were generated using multiple unpaired Welch’s t-tests with log_2_ transformed gene counts. Left and right testicular masses were averaged before statistical analysis. Statistical significance for gene fold changes in Table 1 calculated using multiple unpaired t-test on normalized values. Statistical analyses were done using R version 4.6.0 and Prism 11 software (Graphpad Prism version 11.0.0), p-value lower than 0.05 was considered significant.

**Table 1.**
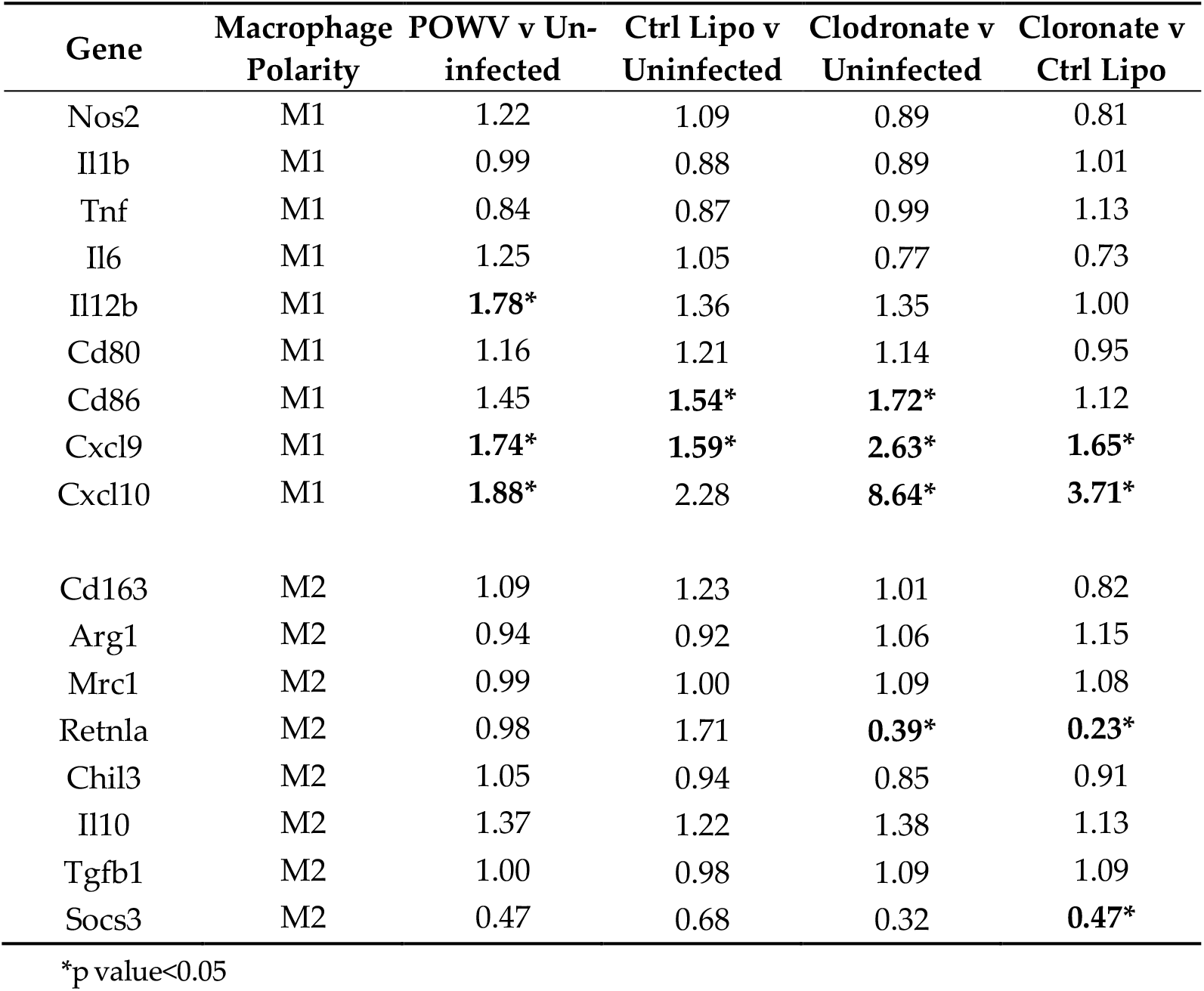
Fold change in normalized expression of M1-and M2-associated gene expression.

## 3. Results

### 3.1. Powassan virus replicates efficiently in primary human Sertoli cells

To determine whether POWV is capable of infecting testes, we first assessed viral replication kinetics in primary human Sertoli cells (HSerCs) as an *in vitro* test for viral tropism. Sertoli cells are a long-lived key testicular cell type responsible for maintaining the blood-testis barrier, thereby shielding the immune privileged niche of the seminiferous tubule lumen from antigen. POWV growth kinetics in BHKsa cells were performed to establish a baseline in a permissive cell type (Figure 1A). Viral titer reached 9 log_10_ PFU/mL by 2–3 days post-infection (dpi) before tapering in titer after 3dpi coinciding with observed cytopathic effect. Similar to BHKsa cells, HSerCs infected with POWV at MOI of 1 and 10 supported robust viral replication, with peak viral titers exceeding 9 log_10_ PFU/mL by 2 days post-infection (Figure 1B). Unlike BHKsa cells, HSerCs maintained high virus titers until 5dpi, and there was a lack of observed cytopathic effect. For comparison ZIKV infection of HSerCs also resulted in productive infection (Figure 1C), included as comparison due to its known tropism for testes^23^.

**Figure 1.**
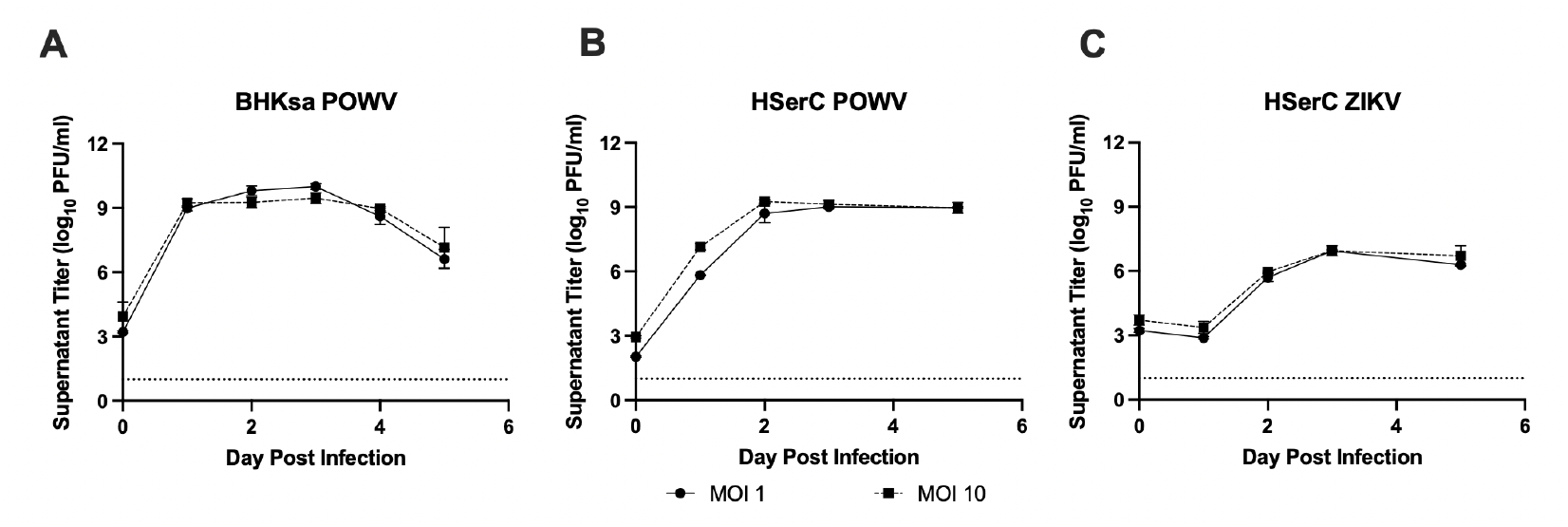
Powassan virus replicates efficiently in primary human Sertoli cells and BHKsa cells. Monolayers of BHKsa cells (A) and primary human Sertoli cells (HSerCs) (B) were infected with POWV or HSerC with ZIKV (C) at multiplicities of infection (MOI) of 1 (circle) or 10 (square). Supernatants were collected daily from 0 to 5 days post-infection (dpi), and viral titers were quantified by plaque assay and expressed as log_10_ PFU/mL. Cell monolayers were observed for cytopathic effect compared to uninfected controls each day. Data shown are representative of three independent experiments. Error bars indicate standard deviation. The dashed line represents the limit of detection (LOD).

### 3.2. POWV infects the male reproductive tract in mice during early infection

To evaluate whether POWV infects the male reproductive tract *in vivo*, adult male BALB/c mice were infected by either the intraperitoneal (IP) or intradermal (ID) route with POWV (doses ranged from 1.9-2.7 log_10_ PFU/mouse), and blood and tissues were collected at defined timepoints post-infection (Figure 2A). Both lineage I (POWV, LB strain) and lineage II (DTV, Spooner strain) were used to rule out a lineage-specific phenotype. Both viruses reached a peak viremia by 3dpi; POWV fell below the limit of detection in both IP and ID route by 5dpi, while DTV remained detectable in blood in 4/5 animals between 2.3-3.6 log_10_ PFU/mL (Figure 2B). Infectious virus was recovered in both testes (Figures 2C and D) and both epididymides (Figures 2F and 2G) of all animals included in the study. POWV was consistently detected as early as 3dpi, with viral loads continuing through 5dpi. Viral titers in testes reached 3.2-6.1 log_10_ PFU/gram tissue by 5dpi. Though testicular infection was ubiquitous from 3-5dpi, by 10dpi only 3/6 (50%) of testes from 2/3 (66%) mice harbored detectable virus. Brain tissue was collected as a positive control of viral dissemination as well as a point of comparison to previously published POWV infection kinetics in BALB/c mice since the central nervous system is a canonical target of POWV (Figure 2H). These data demonstrate early neuroinvasion by 3dpi with more aggressive infection following at 10dpi. Notably, POWV and DTV infection of mouse testes was associated with an increase in testis mass at 5dpi (Figure 2E, supplemental figure 1). Together, these data show POWV infection in the male genital tract regardless of route of infection or viral lineage, so the LP stain given IP was chosen for subsequent studies.

**Figure 2.**
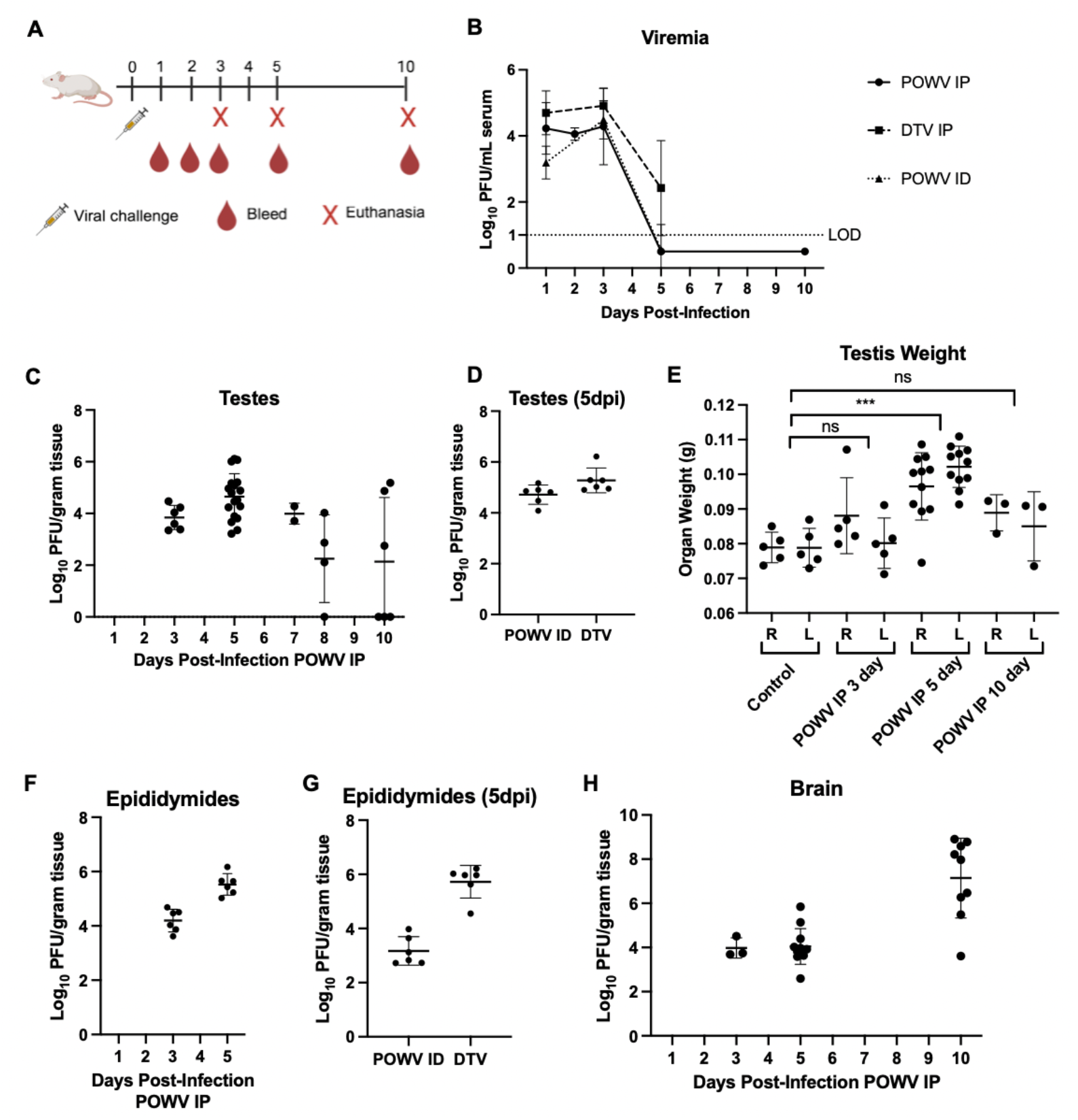
POWV infects the male reproductive tract in mice. Adult male BALB/c mice (n=5) were infected with POWV with 2.08-2.74 _log10_ PFU POWV or 1.98 _log10_ PFU DTV. Schematic diagram of infections and sampling **(A)**. Viremia serum titers are expressed as log_10_ PFU/mL (**B**). Testes, epididymides, and brain collected after euthanasia and titrated to isolate infectious virus (**C**,**D**,**F-H**). Each dot represents an individual organ. Masses of individual mouse testes were measured and separated by right (R) versus left (L) testis, with statistical analysis comparing each condition against control, comparing testes’ weight averages per animal (**E**). Error bars represent standard deviation. IP=intraperitoneal challenge, ID=intradermal challenge, LOD=limit of detection. Ns=not significant, *p<0.05, **p<0.005, ***p<0.0005.

### 3.3. POWV localizes to seminiferous tubules without overt histopathological damage

From each mouse group, between 1/3 to 2/5 of mice testes were paraffin embedded and sectioned for histopathological analysis. Hematoxylin and eosin (H&E)-stained sections of testes from infected mice at 5 and 10dpi were reviewed by a blinded board-certified pathologist and were reported to show no difference in mean seminiferous tubule diameter (mean 0.22 mm diameter in control, mean 0.23 mm diameter in POWV-infected), spermatogenesis potential (<50% of all seminiferous tubules analyzed), or total population of Leydig cells (mean 31.6 Leydig cells/10 high power fields 400x in control, mean 31.1 Leydig cells/10 high power fields 400x in POWV-infected) compared to uninfected controls (Figure 3A–C). There was a lack of observed inflammation nor tubular hyalinization. Immunofluorescence analysis revealed the presence of viral antigen within the seminiferous tubules of infected mice testes at 5dpi (Figure 3D–E).

**Figure 3.**
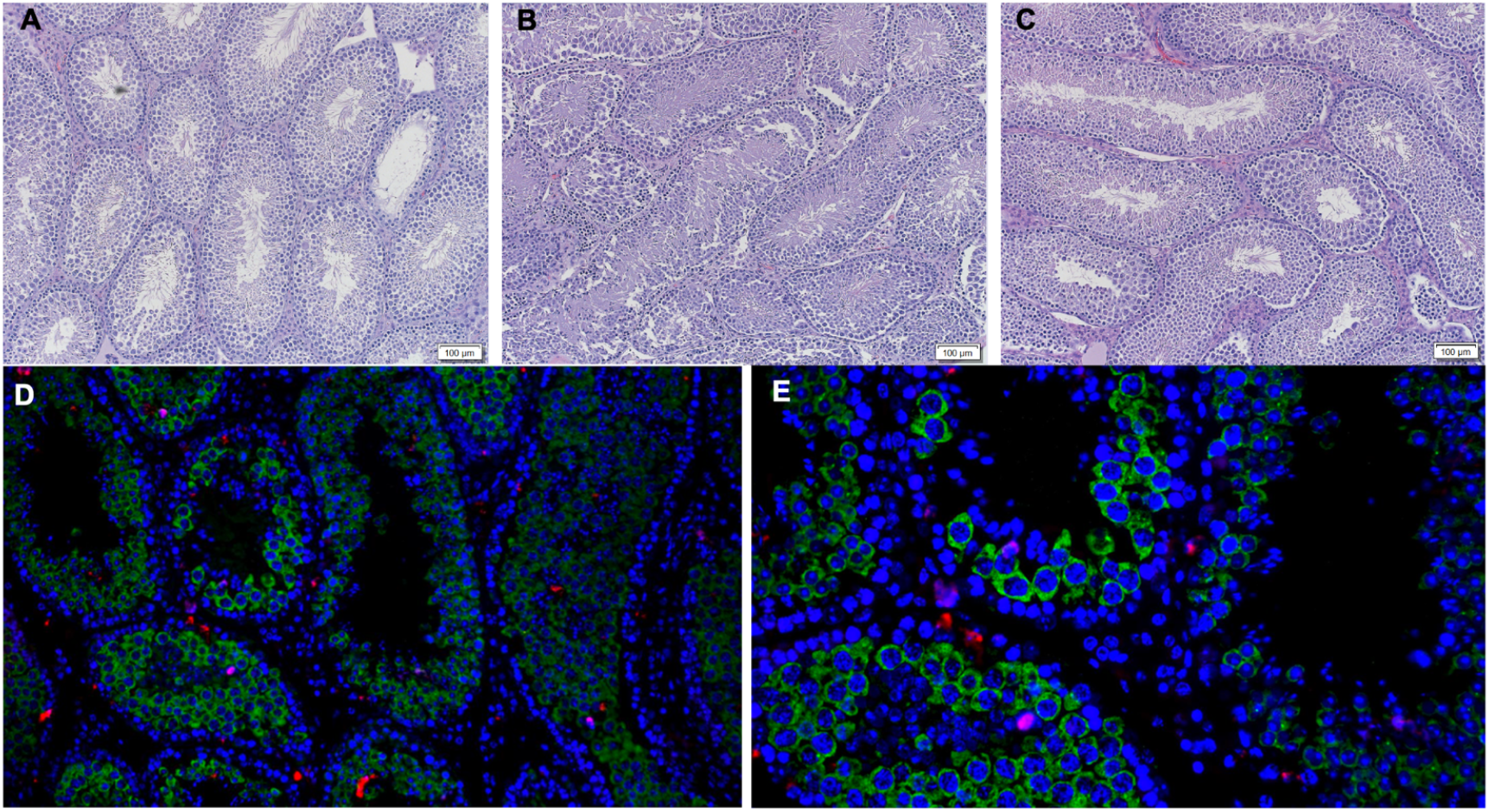
Histological Images of testes from POWV-infected mice. Hematoxylin and eosin (H&E)-stained sections of testes from uninfected control mice **(A)** or POWV-infected mice at 5dpi **(B)** and 10dpi **(C)**. Images shown are representative of all animals examined. Scale bar = 100 µm. Immuno-fluorescence analysis of testis sections from POWV-infected BALB/c mice at 5dpi **(D, E)**. Sections were stained with DAPI (blue) to label nuclei, anti-DDX4 (green), and 4G2 (red). Representative images are shown at 200X **(D)** and 400X **(E)** magnification.

### 3.4. Phagocytic cell depletion enhances systemic viral burden and increases infection of the male reproductive tract

Given that POWV demonstrated tropism for the male reproductive tract, we wanted to evaluate if macrophages facilitated POWV trafficking to the testes contributing to testicular infection. To investigate the role of phagocytic cells in controlling POWV infection, mice were treated with clodronate-containing liposomes to deplete phagocytic cells (Figure 4A). We first performed a clodronate depletion experiment to assess the efficacy of the drug. Percent knockdown in the spleen and the testes was determined using two cohorts of three male BALB/c mice, one cohort treated with clodronate and one cohort treated with control liposomes. After two treatments at 24- and 48-hours prior to euthanasia, spleens and testes of animals were collected and analyzed via flow cytometry. Splenic macrophages depleted 93% after treatment and testicular macrophages were found to increase (Supplemental Figure 2).

**Figure 4.**
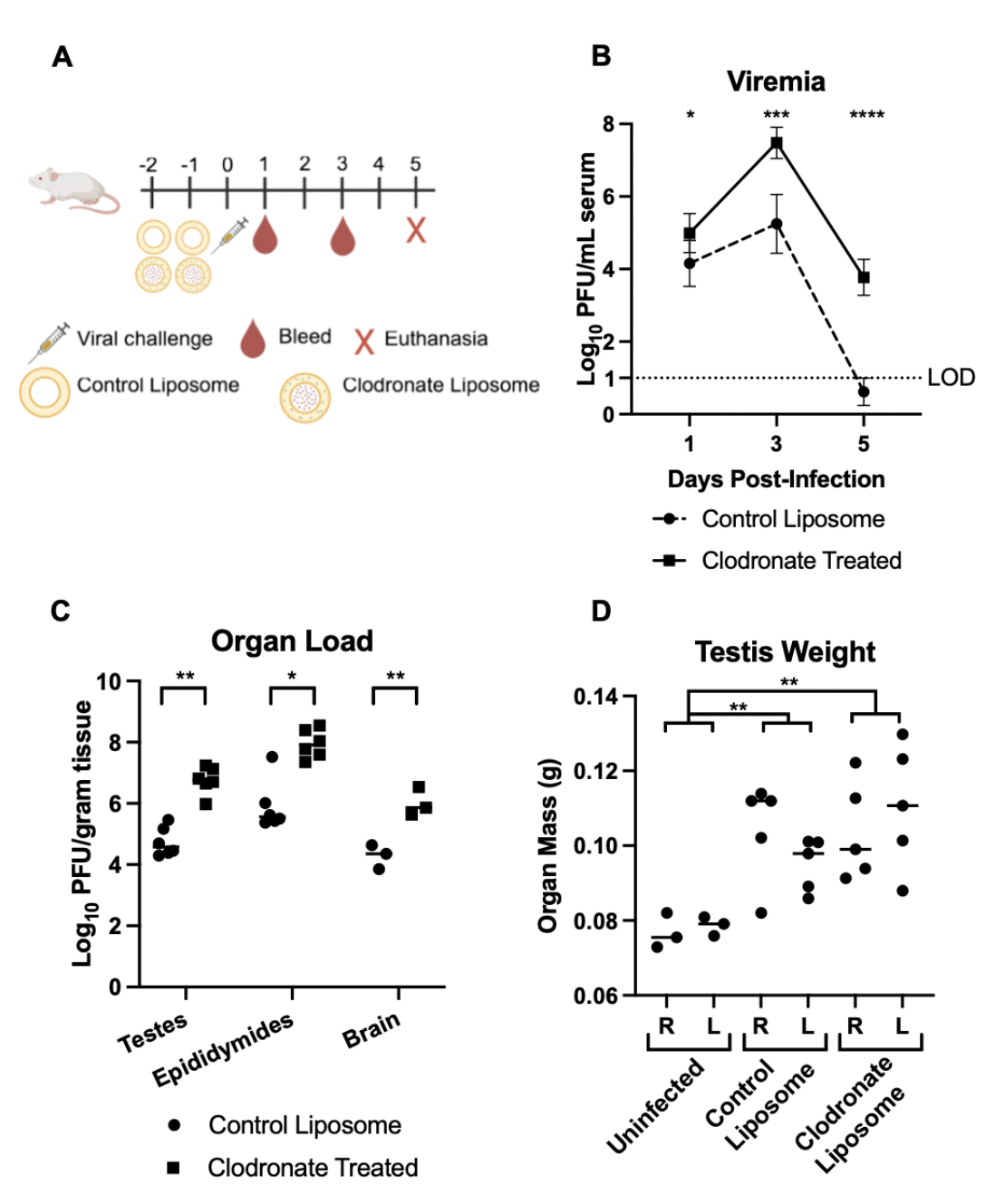
Phagocytic cell depletion enhances systemic Powassan virus burden across multiple organs. Adult male BALB/c mice (n=5) were treated with clodronate-containing liposomes or control liposomes prior to infection with POWV (A). Mice were challenged IP with 2.08 log_10_ PFU POWV. Viremia serum titers are expressed as log_10_ PFU/mL (B). Organs of the MGT were collected on 5dpi (C). Mass of left and right testes on 5dpi (D). Viral titers were quantified by plaque assay and expressed as log_10_ PFU/mL. Limit of detection (LOD) indicated by dashed line. Statistical significance for graph B was determined through a two-way ANOVA, and statistical difference for graphs C and D were determined through paired student’s t test (*p < 0.05, **p < 0.005, ***p < 0.0005, ****p < 0.00005; ns, not significant).

Phagocytic cell depletion resulted in significantly increased and prolonged viremia compared to control liposome-treated animals (Figure 4B). Consistent with this increase in viremia, viral titers in testes, epididymides, and brain were significantly elevated in phagocytic cell-depleted mice (Figure 4C). Additionally, testes from infected mice exhibited increased mass compared to uninfected controls (Figure 4D).

### 3.5. POWV and phagocytic cell depletion are associated with transcriptional shifts in testicular tissue

To characterize the local immune response associated with POWV testicular infection and determine the impact of phagocytic-cell depletion, we performed NanoString transcriptional profiling on whole testis homogenates collected at 5dpi from PBS-perfused mice. Principal component analysis plot assesses transcriptomic cluster behavior among groups (Supplemental figure 3) and differential gene expression analysis is visualized through volcano plots comparing either POWV-infected vs uninfected mice (Figure 5A) or clodronate-treated POWV-infected vs control liposome-treated POWV-infected mice (Figure 5B). PCA plots show biological variation between mice in the study, with experimental groups mostly separating. Overlap of POWV-infected mice with POWV-infected mice challenged with control liposomes in the PCA analysis indicates that the liposomes themselves did not substantially alter the testicular transcriptomic profile. POWV infection was associated with transcriptional induction of antiviral and inflammatory genes, including increased expression of interferon-stimulated genes such as *Irf7* and *Isg15*, as well as inflammatory chemokines including *Cxcl10* and *Ccl8*. Genes associated with myeloid immune activation and inflammatory innate immune responses, including *Fcgr1, Fcgr4, S100a9*, and *Ly6c1*, were also elevated in infected animals. Increased expression of immune regulation and tissue remodeling genes including *Il1b* and *Elane* was also observed. Compared with infected control liposome-treated mice, clodronate-treated animals demonstrated further increased expression of several inflammatory and antiviral response genes, including *Cxcl10, Fcgr1*, and *S100a9*.

**Figure 5.**
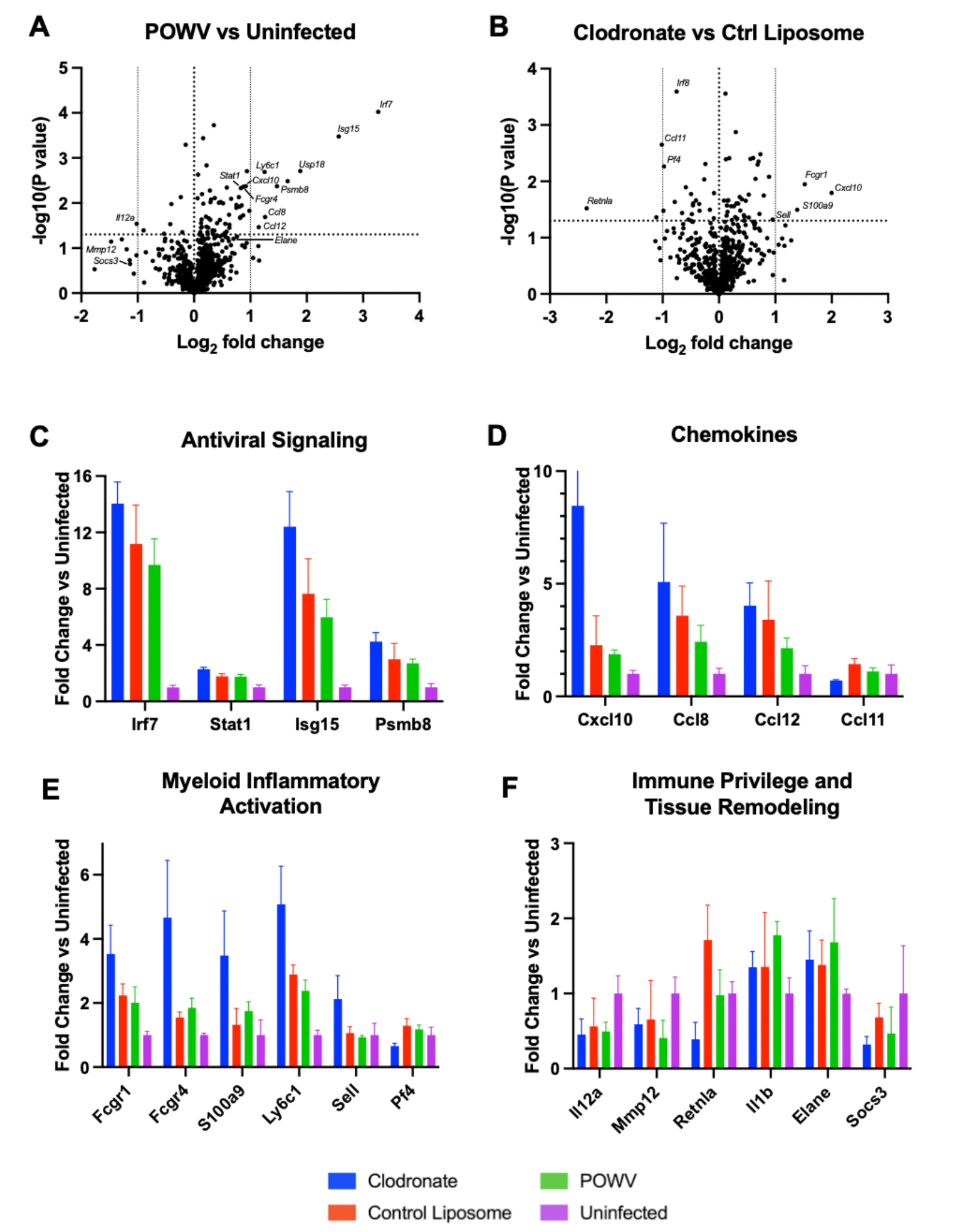
POWV infection induces inflammatory and antiviral gene expression in the testes, which is enhanced following phagocytic cell depletion. NanoString gene expression analysis was performed on testis organ homogenate from mice 5dpi using mouse Myeloid Innate Immunology V2 and Mouse Immunology V2 panels. Groups include clodronate liposome treated POWV-infected, control liposome treated POWV-infected, POWV-infected, and PBS-treated mice. Volcano plots (A,B) were generated using log2 transformed values using unpaired Welch’s t-tests. Fold change of log_2_ transformed normalized read values from selected genes associated with antiviral signaling (C) (*Irf7, Stat1, Isg15*, and *Psmb8*), inflammatory cytokine responses (D) (*Cxcl10, Ccl8, Ccl12*, and *Ccl11*), myeloid associated immune activation (E) (*Fcgr1, Fcgr4, S100a9, Ly6c1, Sell*, and *Pt4*), and immune privilege and tissue remodeling pathways (F) (*Il12a, Mmp12, Retnla, l1b, Elane*, and *Socs3Usp18*). Error bars represent standard deviation.

## 4. Discussion

In this study, we report that POWV consistently infects the male reproductive tract of mice during the acute phase of infection. Infectious virus was recovered from each analyzed testis and epididymis collected at 3- and 5dpi regardless of viral lineage or challenge route. Immunofluorescence imaging revealed POWV envelope protein localized within the seminiferous tubules, implying that POWV may be capable of breaching the blood-testis barrier (BTB). Notably, despite the high testicular titer and presence of viral antigen, there was a lack of observed acute histopathological events. In some mouse models, especially at post-acute timepoints, ZIKV infection of mice can be associated with severe pathological damage including tubular necrosis and impaired spermatogenesis ^13,23^. The absence of overt pathology and lack of disrupted testicular architecture may be a result of the relatively early time points of our study, and further analysis is required to determine if testicular damage may be an outcome beyond our study’s scope of testicular POWV infection.

POWV is primarily recognized as a neuroinvasive virus, and its tropism for the central nervous system has been well supported by clinical and animal model data. A 2016 clinical case of a 63-year-old man who succumbed to POWV disease challenged the current understanding of disease progression by initially presenting as testicular pain^6^. Post-mortem analysis found extensive neuropathological findings; however, the postmortem presence of POWV antigen in the patient’s seminiferous tubules provided the first evidence of potential testicular infection by this virus. The patient was taking rituximab antibody therapy, which obscures whether the presence of testicular viral antigen was evidence of a previously unknown tropism or a rare disease outcome as a consequence of antibody therapy. Rituximab depletes B cell populations, which can impair humoral immunity response to viral infections leading to atypical outcomes of viral disease^24^. Impaired humoral immunity can alter viral disease severity and result in altered viral tropism, including prolonged enteroviral meningoencephalitis, severe neuroinvasive West Nile virus, and prolonged SARS-CoV-2 infection^24^. Despite this, the confirmation of even one case of testicular infection, albeit suspect, warrants further study given the paucity of human cases.

Our study reports POWV testicular infection in mice between 3 and 10dpi. A noteworthy feature of viral testicular infection is the potential to establish a persistent infection, similar to what is reported in ZIKV and EBOV infections^13,25,26^. Our study reported 3/6 testes in 2/3 mice positive for POWV infection at 10dpi, without insight into the viral dynamics that may have resulted at later time points. *Raney et. al*. reported high testicular titer at 28dpi in 1/5 mice tested, suggesting that long-term testicular POWV infection in mice is a potential outcome^18^.

Consistent with the clinical outcome of orchiepididymitis in the previously discussed patient, POWV-infected mice in our study were associated with increased testicular mass at 5dpi. This finding is contrasted with ZIKV infection, which causes testicular atrophy and reduced testicular mass in murine models prior to 10dpi^27^. ZIKV mouse models are typically interferon signaling impaired, which can exacerbate disease outcomes, preventing a direct comparison from our immunocompetent mouse model. These findings suggest infected mice are experiencing testicular edema and swelling due to a proinflammatory response at the testes during POWV infection. This phenotype is consistent with our gene expression analysis results that confirm a robust antiviral and pro-inflammatory host response at the testes.

Due to macrophages implicated in driving ZIKV testicular infection in mice through a “Trojan horse”-like mechanism, we sought to determine if POWV uses a similar mechanism^21^. Interestingly, we found that after phagocytic-cell depletion, POWV viral load in the testes was significantly increased, indicating that POWV most likely reaches the testes independent of infected macrophages. In addition, depletion of phagocytic cells was associated with significantly higher and sustained POWV viremia. Phagocytic cells play key roles in mediating both pro- and anti-inflammatory pathways especially at the testis, where their anti-inflammatory phenotype contributes to maintaining immune privilege^28^. Our study reflects this dynamic with shifting transcriptional levels of M1 and M2-associated genes present in the experimental conditions evaluated (Table 1). Notably, when phagocytic cells were depleted in the testes, the anti-inflammatory M2-associated gene *Retnla* was significantly downregulated. The loss of this M2-associated gene and the upregulation of M1-associated genes during phagocytic-cell depletion reflect the baseline anti-inflammatory environment of the testes. Upregulation of M1 associated genes during infection reflect the induction of a proinflammatory response at the testis. Our data suggest that phagocytic cells play a protective role in clearing viremia and controlling testicular POWV infection. This finding challenged expectations, as monocyte-derived cells are often implicated in driving arbovirus dissemination.

Gene expression analysis of perfused testes harvested 5dpi indicated a strong antiviral and inflammatory response associated with POWV infection. Interferon-stimulated genes, antigen presentation pathways, chemokine signaling, and myeloid activation markers were consistently upregulated transcriptionally. Among the highest upregulated genes, *Irf7* and *Cxcl10* represent pro-inflammatory transcriptional signal in the testes, and *Fcgr1, Fcgr4, Ctss*, and *Ccl8* transcriptional upregulation imply recruitment and activation of innate immune cell populations. These indicate a strong pro-inflammatory response induced in the testes, an immune privileged space, likely contributing to the observed increase in testicular mass despite overt signs of inflammation in the H&E stained histology slides. In addition to transcripts associated with classical pro-inflammatory pathways, genes responsible for epithelial barrier integrity and immune privilege regulation are of particular interest in the testes. Downregulation of *Socs3*, an anti-inflammatory gene linked to immunoregulatory signaling in the testes, could indicate potentially impaired immune privilege maintenance^30^. These factors could contribute to disruption of the delicate immune homeostatis within the immune privileged environment of the testes. Upregulation of *Il1b* may imply BTB involvement, as *Il1b* is associated with epithelial tissue disruption, and interleukin-1 family proteins regulate BTB restructuring^31,32^. Furthermore, *Elane*, another upregulated cytokine, is associated with disruption of epithelial barriers; relationships between the BTB and this cytokine have yet to be directly evaluated, but *Elane*-mediated disruption of other epithelial barriers is through cleavage of the same tight-junction proteins that contribute to BTB strength^33,34^. These transcriptional insights may imply potential mechanisms for POWV-mediated BTB disruption; however, further analysis is required to address this directly.

As phagocytic cell depletion was confirmed in a control cohort at time of infection, the increased and prolonged viremia observed was most likely due to an impaired innate immune response. The high testicular organ viral load and transcriptional response is most likely a downstream response to the robust early viral replication due to the loss of a typical antiviral response, similar to what has been observed with west Nile virus infection of macrophage-depleted mice^29^. Infected testes at 5dpi in clodronate-treated mice therefore show potential compensatory induction of *Cxcl10, Ccl2, Ccl5*, and *Fcgr4* upregulated compared to liposome control-treated mice. These reflect the host response to an increase viral burden. Due to macrophage depletion only being confirmed at day 0, our data do not evaluate the level of macrophage depletion in the testes at 5dpi, and transcriptomic data show myeloid-associated genes in testes at 5dpi. Previous depletion studies show that recovery of testicular macrophages can take over 30 days after depletion^35^. These transcriptional findings are consistent with enhanced immune activation in the testes following phagocytic-cell depletion and parallel the increased viremia and higher testicular viral burden observed in clodronate-treated animals.

Though infection of testes and epididymides was consistently observed in mice, it remains unclear the translatability of our findings to clinical cases. Currently, there have been no reported cases of POWV resulting from human-to-human transmission, and sexual transmission would represent a major departure from the current understanding of POWV epidemiology. We found consistently high testicular POWV viral load in testes at early timepoints prior to presentation of clinical disease manifestations in mice.

We report consistent testicular infection in mice by POWV; however, the potential for POWV to spread sexually requires many unexplored biological events such as shedding in the seminal fluid and a susceptible recipient host. In many cases, viral infection of testes does not correspond to substantial shedding in seminal fluid. For ZIKV, testicular infection is associated with viral shedding in semen resulting in potential sexual transmission. Spondweni virus, an *orthoflavivirus* closely related to ZIKV, reaches similar viral organ loads in the male reproductive tract to ZIKV. Unlike ZIKV, Spondweni virus shedding in semen is a significantly rarer event^36^. Japanese encephalitis virus (JEV) is detected in testes and semen of pigs and boars; however, this does not translate to humans as sexual transmission is not considered a viable route of transmission for JEV^37^. Ultimately, whether POWV sheds in the semen, is capable of transmitting sexually, or if testicular infection is a regular outcome of human disease is unknown and requires further analysis to confirm.

## Supporting information

https://doi.org/10.5281/zenodo.21417492

## Supplementary Materials

The following supporting information can be downloaded at: https://www.mdpi.com/article/doi/s1, Figure S1: title; Table S1: title; Video S1: title.

## Author Contributions

Conceptualization, methodology, validation, formal analysis E.E.H.S, A.N.F, and S.L.R.; investigation, E.E.H.S, E.A, E. E.; resources, A.N.F. and S.L.R.; writing—original draft preparation, review, and editing E.E.H.S. A.N.F, and S.L.R.; supervision, A.N.F. and S.L.R. All authors have read and agreed to the published version of the manuscript.

## Institutional Review Board Statement

The animal study protocol was approved by IACUC protocol 2507034 and performed with guidelines of the Association for Assessment and Accreditation of Laboratory Animal Care (AALAC) International.

## Data Availability Statement

All raw data described in this manuscript including Supplamentary data are available for access through Zenodo Data DOI https://doi.org/10.5281/zenodo.21417492

## Acknowledgments

Viruses used in this study were provided by the World Reference Center for Emerging Viruses and Arboviruses at the University of Texas Medical Branch. Graphs were generated using Graphpad Prism version 11.0.1. The authors would like to thank Dr. David H. Walker in analyzing histology slides of mouse testis, Meredith Weglarz for assistance in analyzing samples via Flow Cytometry, Dr. Marielena Vogel Saivish for experimental support.

## Abbreviations

The following abbreviations are used in this manuscript:

POWV: Powassan virus
DTV: Deer tick virus
ZIKV: Zika virus
dpi: days post-infection
PFU: plaque forming units
FFU: focus forming units
HSerCs: primary human Sertoli cells
BTB: blood-testis barrier
ABSL3: animal biosafety level 3
MOI: multiplicity of infection
IP: intraperitoneal
ID: intradermal
H&E: hematoxylin & eosin
EBOV: Ebola virus
JEV: Japanese encephalitis virus

