## Supplementary material for "Powassan virus infects mouse testes and reveals a protective role for phagocytic cells": https://doi.org/10.5281/zenodo.21417492

### Supplemental Figures

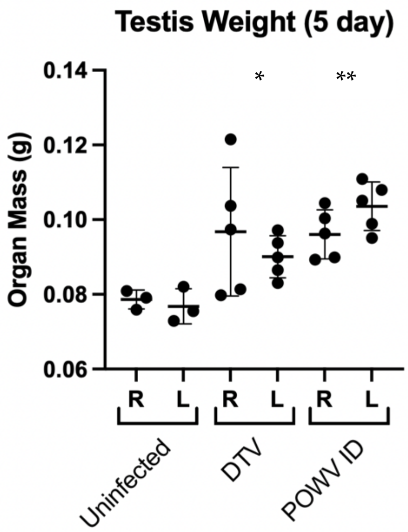


**Supplemental Figure 1**: Adult male BALB/c mice (n=5) were infected with 2.08-2.74 $\log_{10}$ plaque forming units (PFU) POWV or 1.98 $\log_{10}$ PFU DTV. Individual mouse testes weighed and separated by right (R) versus left (L) testis, with statistical analysis comparing each condition against control via student's t test, comparing testes' weight averages per animal.

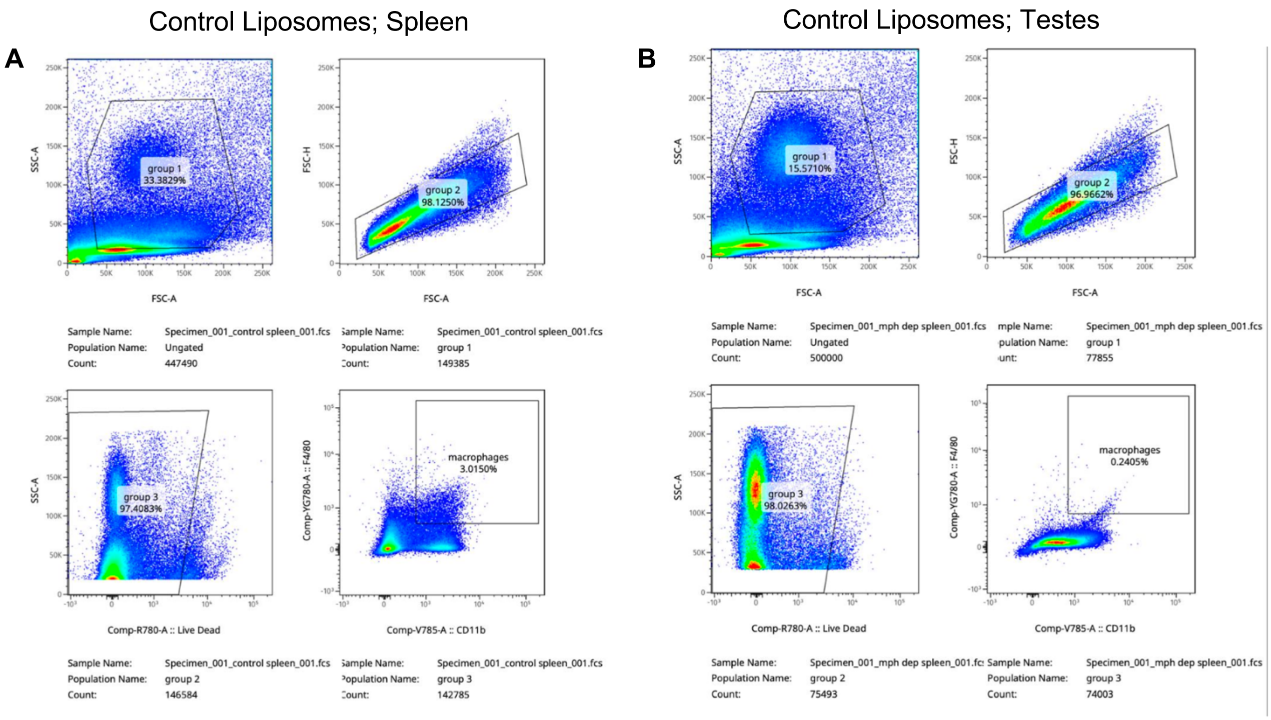

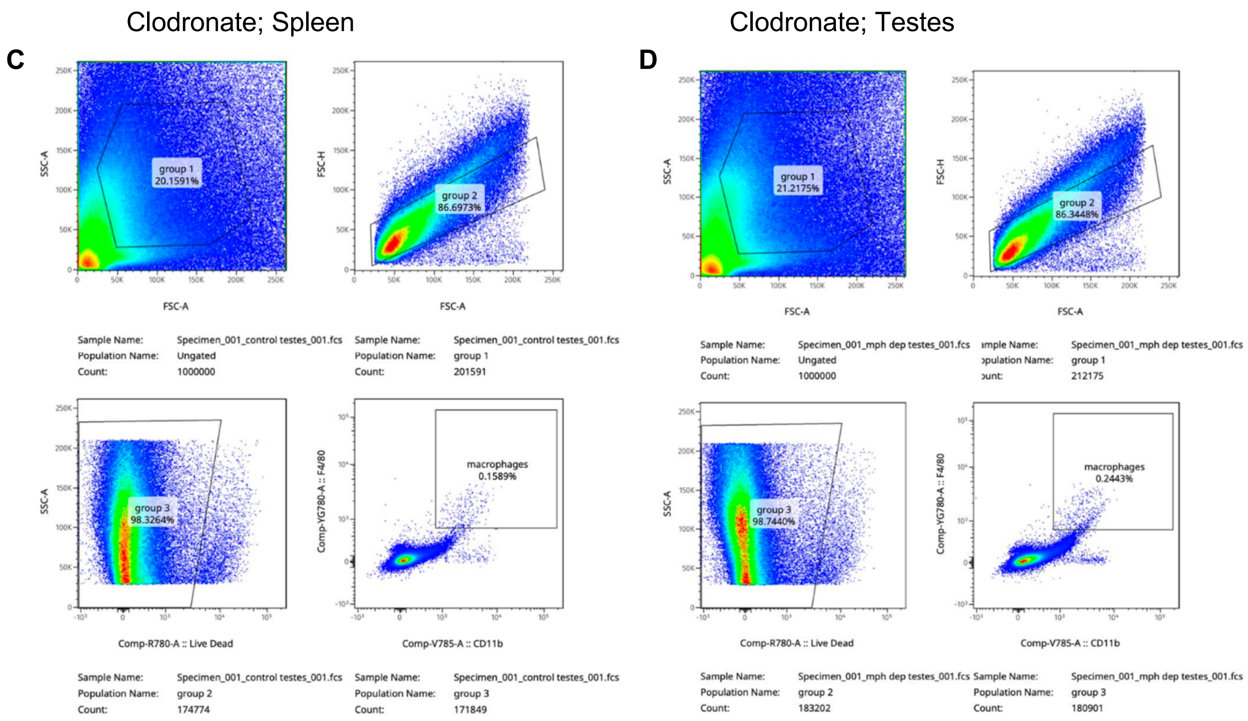


**Supplemental Figure 2:** Clodronate liposome result in depletion of phagocytic cells in spleen but not testes 1-2 days post-injection. BALB/c mice (n=3/treatment) were injected IP with either clodronate containing liposomes or control liposomes 24- and 48-hours prior to euthanasia and organ collection. Animal spleens and testes were collected and processed for flow cytometry as detailed in material and methods section. Figures above show gating strategy for clodronate-treated animals (A,B) and control liposome-treated animals (C,D). Live singlet cells were sequentially gated and determined to be macrophages through expression of both Cdllb and F4/80. Gating for antibodies was determined through use of compensation controls. Percentages shown are representative of initial parent gate. Data presented show results from one animal and are representative of three tested mice.

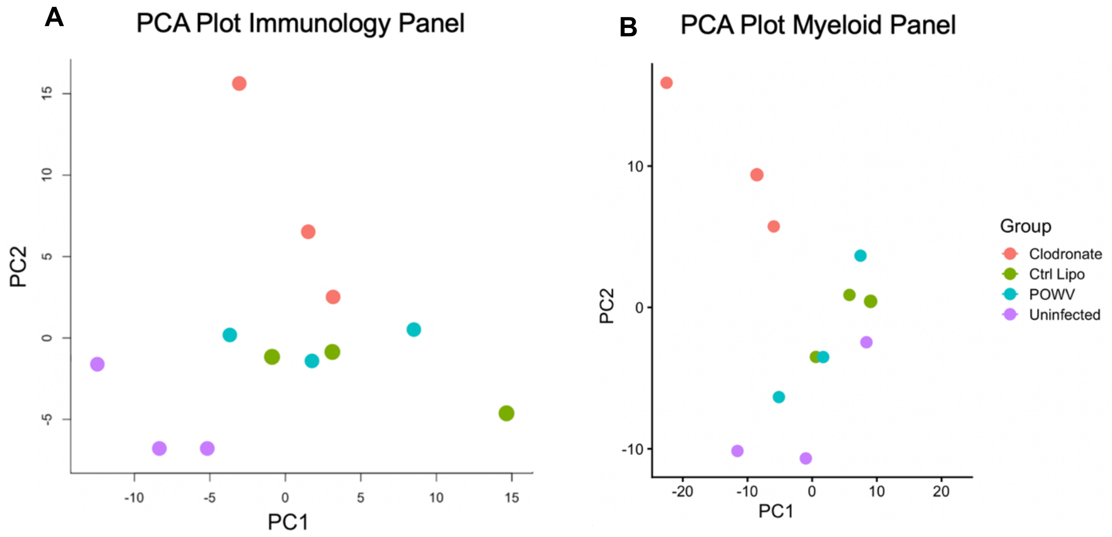


**Supplemental Figure 3:** Principal component analysis (PCA) of gene expression analysis results for the immunology (A) and myeloid (B) panels. PCA was performed using $log_2$-transformed normalized gene expression data. Each point represents an individual animal, with testes combined.
